# Unveiling diffusion pattern and structural impact of the most invasive SARS-CoV-2 spike mutation

**DOI:** 10.1101/2020.05.14.095620

**Authors:** Emiliano Trucchi, Paolo Gratton, Fabrizio Mafessoni, Stefano Motta, Francesco Cicconardi, Giorgio Bertorelle, Ilda D’Annessa, Daniele Di Marino

## Abstract

Starting in Wuhan, China, SARS-CoV-2 epidemics quickly propagated worldwide in less than three months, geographically sorting genomic variants in newly established propagules of infections. Stochasticity in transmission within and between countries and/or actual advantage in virus transmissibility could explain the high frequency reached by some genomic variants during the course of the outbreak.

Using a suite of statistical, population genetics, and theoretical approaches, we show that the globally most represented spike protein variant (*i.e.*, the G clade, A → G nucleotide change at genomic position 23,403; D → G amino acid change at spike protein position 614) *i)* underwent a significant demographic expansion in most countries not explained by stochastic effects or enhanced pathogenicity; *ii)* affects the spike S1/S2 furin-like site increasing its conformational plasticity (short range effect), and *iii)* modifies the internal motion of the receptor-binding domain affecting its cross-connection with other functional domains (long-range effect).

Our study unambiguously links the spread of the G614 with a non-random process, and we hypothesize that this process is related to the selective advantage produced by a specific structural modification of the spike protein. We conclude that the different conformation of the S1/S2 proteolytic site is at the basis of the higher transmission rate of this invasive SARS-CoV-2 variant, and provide structural information to guide the design of selective and efficient drugs.

## INTRODUCTION

After appearing in Wuhan, China, in late 2019, SARS-CoV-2 (Zhou et al 2020, WHO 2020a), a highly contagious (Liu et al 2020, D’Arienzo and Coniglio 2020) coronavirus (CoV) causing severe acute respiratory syndrome COVID-19, spread worldwide and rapidly emerged as a dramatic pandemic, officially acknowledged on March 11 2020 (WHO 2020a). As of the 12th of May 2020, about 290,000 deaths related to COVID-19 have been recorded, mainly in the US, UK, Italy, Spain and France, while the global number of infected people far exceeded four million (WHO 2020b).

All CoVs encode a spike glycoprotein presented on the surface of the viral particle as a trimer where each monomer is composed by two subunits (S1 and S2). After cleavage by host proteases, S1 and S2 subunits remain non-covalently bonded. The S1 subunit contains a N-terminal domain (NTD), a receptor-binding domain (RBD) that drives host cell tropism and a C-terminal domain further subdivided in domains SD1 and SD2, while the S2 subunit mainly consists of heptad repeat (HR) regions involved in membrane fusion (Belouzard et al 2012, Liu et al 2004, Li 2016; Fig. 1A-B). Spike protein monomers can exist in two main metastable conformations: down, with the RBD tightly packed against the NTD, and up (Song et al 2018; Walls et al 2020; Wrapp et al 2020). The up conformation represents the active form of the spike protein, corresponding to the receptor-accessible configuration (Berry et al 2004, Pak et al 2009, Walls et al 2017, Wrapp et al 2020). In the cryo-EM structure of the spike protein only one monomer is found in the up configuration and expected to contact the human angiotensin-converting enzyme 2 (ACE2; Walls et al 2020, Zhou et al 2020), mediating host cell invasion.

**Figure 1.**
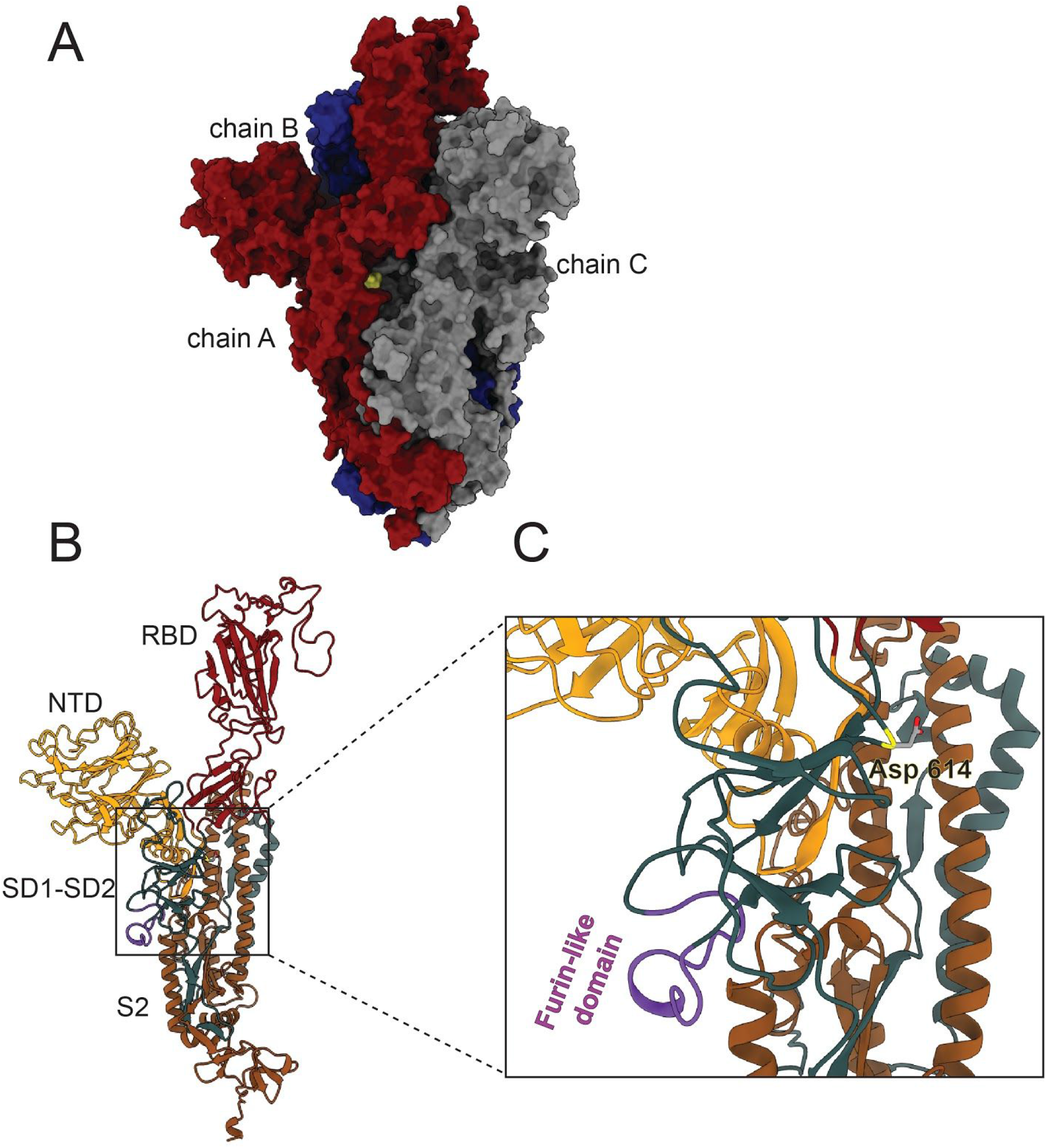
Three-dimensional structure of Spike protein. **A.** Molecular surface of the SARS-CoV-2 spike protein. Chain A in red is in the up conformation, while chains B and C in blue and grey respectively are in the down conformation. The site of the D614G substitution is marked in yellow. **B.** Domain subdivision of the monomer. NTD: N-terminal of S1 subunit (dark yellow); RBD: receptor-binding domain (dark red); SD1-SD2: C-terminal region of S1 subunit (petrol green) harbouring the furin-like domain (purple); S2: heptad repeat 1 (HR1) in the S2 subunit (orange); **C.** Close-up on the region where the D614G variant (Asp 614; yellow) is located. The furin-like domain is highlighted (purple).

Coronavirus spike proteins must be primed before they can be triggered to induce fusion between the viral and the host cellular membranes (Simmons et al 2013, Hoffmann et al 2020a). Priming involves a proteolytic event at a cleavage site (S2’), mediated by the cellular serine protease TMPRSS2, which converts the protein from a fusion-incompetent to a fusion-competent state (White & Whittaker 2016). One of the evolutionary innovations in SARS-CoV-2, which has been suggested to enhance its infectivity (Andersen et al 2020, Coutard et al 2020), is the presence in the spike of a novel and peculiar furin-like cleavage site (S1/S2) in the form of an exposed loop harbouring multiple arginine residues (PRRAR/S amino acid sequence cleaved at the ‘/’ symbol; Walls et al 2020; Wrapp et al 2020; Fig. 1B-C). This site has been observed, though with different amino acid sequences, in distantly related CoVs, like MERS-CoV and HKU1-CoV, but not in those viruses that are the most closely related to SARS-CoV-2 (Coutard et al 2020, Andersen et al 2020, Xiao et al 2020).

The SARS-CoV-2 spike is then activated by a two-step process: a priming cleavage by furin-like protease at the S1/S2 site and an activating TMPRSS2-mediated cleavage at the S2’ site during membrane fusions (Ou et al 2020, Hoffmann et al 2020b). The S1/S2 cleavage site is also involved in cell-cell transmission via syncytium formation in SARS-CoV-2 as well as in MERS-CoV (Qian et al 2013, Ou et al 2020), a more efficient spreading within the host than cell-free aqueous diffusion (Mothes et al 2010). In laboratory experiments, deletion of the S1/S2 motif resulted in a spike protein that was no longer able to induce syncytium formation whereas modification of the S1/S2 motif with a more efficient one (alanine-to-lysine substitution: RRAR -> RRKR) strongly increased syncytium formation potentially enhancing pathogenicity (Hoffmann et al 2020b).

A novel spike variant of the spike was discovered in Bavaria, Germany (EPI_ISL_406862,) and Shanghai, China (EPI_ISL_422425, 416327, 416334) in late January 2020. Only a few weeks later, this variant emerged as the most abundant clade in Europe (Becerra-Flores and Cardozo 2020, Brufsky 2020, Laha et al 2020, Pachetti et al 2020, Chiara et al 2020, GISAID’s EpiFlu Database), and in April it was acknowledged as the prevailing variant worldwide (Korber et al 2020, GISAID’s EpiFlu Database). This SARS-CoV-2 variant is characterized by a nucleotidic transition A → G at the genomic position 23,403, (Wuhan reference genome; Wu et al 2020), changing an aspartic acid at the spike position 614 into a glycine (hereafter G614, whereas the ancestral state is indicated as D614; reported in yellow in Fig. 1).

As G614 prevails in every region where it seeded the epidemic and also where it invaded a region long after the alternative D614, a selective advantage for this novel variant, and/or its linkage group (Pachetti et al 2020), has been suggested (Korber et al 2020, Vasilarou et al 2020). However, stochastic processes such as founder effect (*i.e.*, G614 is frequent because G614 carriers from a common epidemiological source seeded the epidemic in different countries) or shared genetic drift (*i.e.*, common random fluctuations in local populations connected by abundant gene flow) may be responsible for the observed frequency increase of G614 (Chiara et al 2020). In fact, assessing the adaptive significance of any genetic change requires a combined approach based on epidemiological data, population genomics, and a functional explanation of the hypothetical selection advantage.

The molecular bases for the putative G614 selective advantage are not yet understood. The D → G amino acid substitution may affect the activity of a nearby epitope potentially involved in antibody-dependent enhancement, as observed in SARS and MERS-CoV infections (Wang et al 2014, Wan et al 2020). Alternatively, this substitution could affect the molecular interactions in its proximity, facilitating the separation of S1 from S2 when the latter is attached to the cell membrane, or have long-range effects on RBD-ACE2 binding (Korber et al 2020). Detailed analysis of the molecular dynamics of the two alternative spike variants is therefore crucial to identify the molecular changes which could be responsible for the rampaging global diffusion of the G614 and plan specific *in vitro* and *in vivo* assays.

Here, we first demonstrate that the prevalence of the G614 cannot be explained by shared drift and connectivity among epidemics in different geographic areas. Then, we show that the mutation of residue 614 has both short- and long-range effects on the dynamics of the spike protein, affecting both the S1/S2 furin-like cleavage site and the RBD spatial conformation. In particular, in the G614 variant, the multibasic furin-like domain is much more exposed than in the D614 variant, and the position of the cleavage residue R685 more strongly stabilized by the electrostatic interaction with D663, likely facilitating the recognition by the protease. In addition, in the G614 variant, the RBD assumes a more open conformation, which, we propose, may facilitate the interaction with ACE2. The general similarity of observed trends in the relative frequencies of the two alternative variants in the genomic samples and in the actual populations as reconstructed by coalescent-based inference suggest that the G614 molecular features that we identified are more likely boosting transmission rate rather than pathogenicity.

## RESULTS AND DISCUSSION

### Global spread of G614 is not due to random genetic drift

We investigated whether the previously reported frequency increase of the G614 (Korber et al 2020) variant can be attributed to random genetic drift by fitting a generalized mixed model (GLMM) with binomial error structure.

We modeled the probability that a sampled viral genomic sequence has a G at spike amino acid position 614 as a function of time, including random intercepts and slopes for political/administrative units (countries, except for USA States and Chinese provinces, hereafter “territories”) and covariance matrices based on the genetic similarity between viral samples from each pair of territories. The rationale for this model structure is that territories serve as reasonable approximations of genetic drift units (demes) and that *F*_ST_ between two populations can be seen as the correlation of randomly drawn alleles within each population relative to the total. Hence, we used in the model a correlation matrix based on pairwise *F*_ST_ between populations to account for the shared drift among connected territories. Under these assumptions, our fixed effect of sampling day represents the change in frequency that is not explained by genetic drift within territories or is not explained by connectivity among territories.

Our statistical analysis (considering 5,773 sequences from the 24 territories with at least 30 sequences each) confirmed that the observed increase in the relative frequency of the G614 allele is not explained by random drift or connectivity (GLMM: logistic slope per day = 0.096, SE = 0.017, full/null model *LR* = 22.8, *df* = 1, *P* < 0.001). Similar results are obtained when *F*_ST_ is calculated only for sequences carrying the G614 allele (GLMM: logistic slope per day = 0.091, SE = 0.017, full/null model *LR* = 14.0, *df* = 1, *P* < 0.001), in order to account for events of gene flow which introduce this variant at low frequency in one of the two populations, or time is aligned around the midpoint of each population, to control for the correlation between beginning of the epidemics and frequency of G, *i.e.* the intercept (GLMM: logistic slope per day = 0.085, SE = 0.017, full/null model *LR* = 13.4, *df* = 1, *P* < 0.001). Note that the similarity in slope and intercept between population is not correlated with viral population relationships measured as *F*_ST_, further suggesting that migration between viral populations across different territories is not responsible for the widespread increase in the G614 allele over time (Mantel test *p* = 0.96 and *p* = 0.70, for intercept and slope, respectively).

By plotting the random effects for each territory (Fig. 2A), we observed that, although the fitted slope varies among territories, Iceland’s is the only sample to show a decreasing trend in the relative frequency of the G allele (Fig. 2A) over the sampled time window (March, 1-28 2020). At present, we did not collect data allowing us to test whether this idiosyncratic trend in Iceland’s sample can be attributed to any special feature of this country’s epidemic. However, we note that Iceland has recorded one of the world’s lowest fatality rates (ca. 0.5%) estimated through a detailed tracking of the epidemics (covid.is/english).

**Figure 2.**
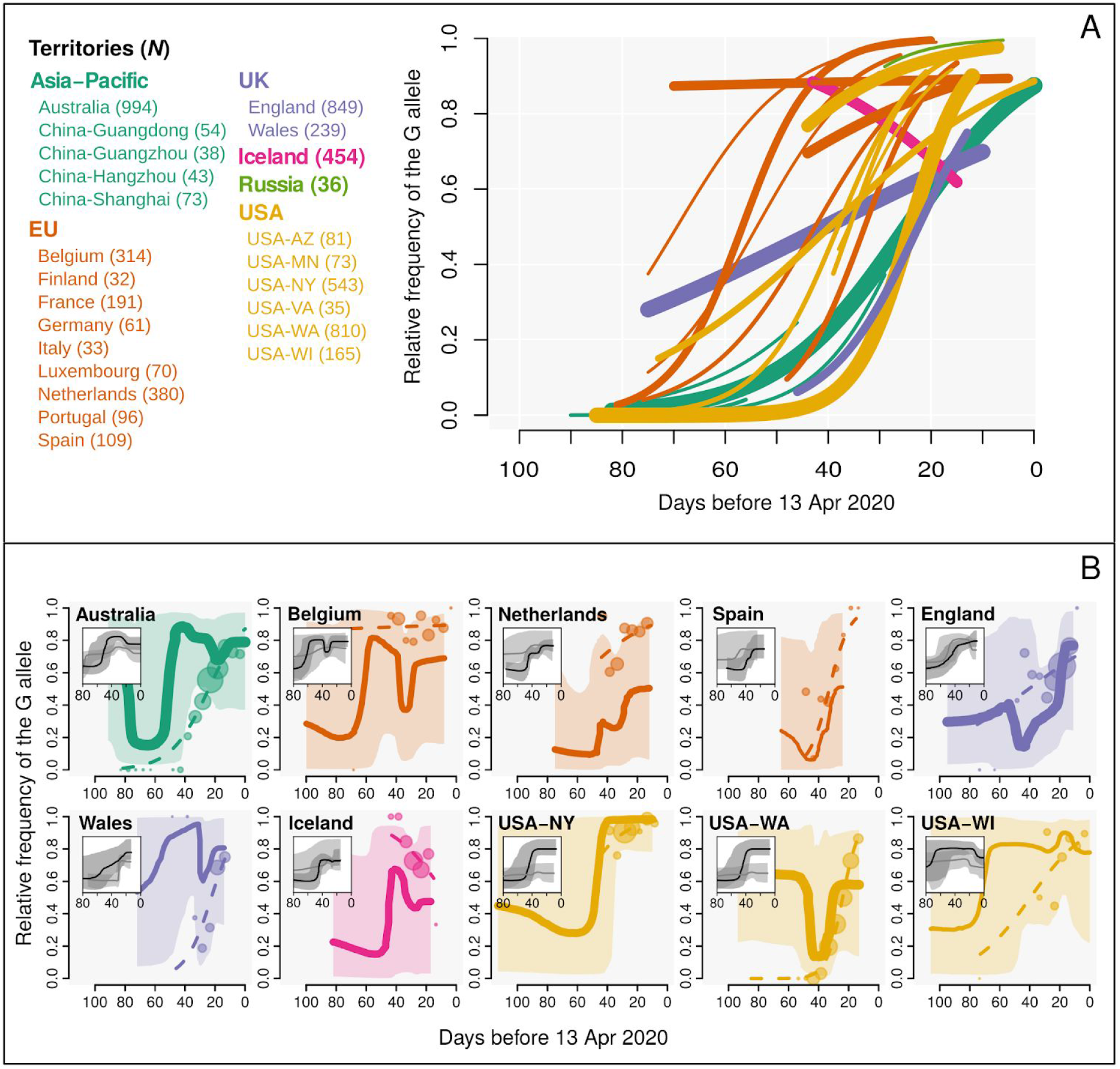
Frequency increase of Spike G614. **A.** GLMM fitted logistic growth for each of 24 territories with > 30 sequenced genomes published as to 13 Apr 2020 (line width proportional to square root of sample size); **B.** Comparison of the frequency of the G614 variant in the genomic samples (circles: raw data aggregated in 5-days intervals, size proportional to square root of sample size; dashed lines: GLMM fit) and in the population as Bayesian Skyline plot ratio: (estimated effective population size - N_e_ - of G614 over the sum of G614 and 614D N_e_; see Methods for details) Solid lines: medians (line width proportional to square root of sample size); shaded areas: 99.75% confidence intervals referring to coalescent-based demographic reconstructions ratio) for the 10 territories with at least 30 sequenced genomes per variant. Demographic reconstructions as inferred by Bayesian Skyline plot for the D614 (gray) and G614 (black) median and 95% CI are shown as insets within each territory panel.

By comparing the logistic slopes from simpler GLMMs without *F*_ST_-based covariance structure, we also observed that the variant G614 and those in the same linkage group (genomic positions 241, 3037 and 14408; Pachetti et al 2020, Zhu et al 2020, Korber et al 2020) show the clearest signal of frequency change across the 24 investigated territories among 15 other variants with minor allele frequency > 0.1 in our dataset (Fig. S1). Among the mutations linked to the 614G variant (genomic position 23,403), the C241T occurs in the short non-coding region before the ORF1a and C3037T is a synonymous change within the ORF1a. On the other hand, C14408T is a non-synonymous change in the *nsp12* gene coding for the RNA-dependent RNA polymerase and could influence replication efficiency or fidelity (Pachetti et al 2020). The structural and functional differences between the ancestral and derived variants of C14408T were not yet thoroughly investigated.

Considering that the genomic samples available through the GISAID EpiFlu Database are mostly coming from hospitalized patients or from patients showing COVID-19 symptoms, the apparent selective advantage of the G614 variant could result from more severe COVID-19 outcomes associated to G614 (*i.e.*, increased pathogenicity), which would increase the prevalence of this variant in the non-random samples of patients seeking treatment. Korber et al (2020) found no association between G614 and symptom severity in a group of patients from a single UK hospital. To generalize this result, supporting enhanced transmission rate over increased pathogenicity in the G614 spread, we used an independent coalescent-based approach to test for the actual dominance of the G614 in different populations (Australia, Belgium, England, Iceland, Netherlands, Spain, Wales, USA-WA, USA-NY, and USA-WI).

By comparing the fitted logistic growth of G614 in each territory with the relative dynamics of the two spike variants in the population as reconstructed by a coalescent-based demographic inference (Bayesian Skyline plots - BSP; Drummond et al 2005), we observe a similar trend of the G614 in the genomic samples as well as in the overall population (Fig. 2B) excluding higher pathogenicity as a potential explanation of the G614 prevalence in the genomic samples and, instead, further supporting the hypothesis of its higher transmission rate. Although the pattern is clear in all territories (*i.e.*, similar general trend in the genomic sample and in the population), in Iceland and Netherlands, the BSP reconstructions show a lower prevalence of the G variant in the population than in the hospitalized samples. We note again that Iceland is the only territory showing a clear decline in the relative frequency of G614 in the sample, which is also tracked by the BSP, but the case of Netherlands (one of the countries contributing the most to virus sequencing from the very beginning of the outbreak; GISAID Database) is more difficult to explain. Even taking into account these differences, we conclude that increased pathogenicity is an unlikely explanation for the overrepresentation of G614 in late COVID-19 samples.

### G614 enhances virus performance making the furin-like cleavage site more accessible

We investigated the selective advantage of G614 by means of molecular dynamics (MD) simulations. The time evolution of D614 and G614 variants was followed for 0.5 µs (for a total of 1 µs) to identify the impact of this mutation on the spike structure. The substitution of an aspartic acid (D) with a glycine (G) at position 614 modifies the whole fluctuation profile of the spike trimers (Fig. S2A), in particular in the region downstream of the mutation (residues 622-642): interestingly, in chain A, where the RBD is in the up conformation, this amino acid substitution increases the fluctuations, while in chains B and C, bearing the RBD in the down conformations, it has an opposite effect, reducing the fluctuations (Fig. S2B). The altered profile of the fluctuation extends farther downstream, toward residues 675-692 (Fig. S2B), comprising the furin-like domain. Of note, the increased mobility of the furin-like domain appears only in chain A with the activated RBD could have a functional meaning.

The increased flexibility in residues 675-692 makes the furin-like domain more solvent exposed, with an increase of the accessible surface (SAS) area of ∼1.0 ±0.7 nm^2^ in G614 as compared to D614 (Fig. 3A) In D614 this domain is tightly bound to the protein surface, whereas the loop samples a larger conformational space in G614. The loop can detach from the protein and protrude towards the solvent, thus increasing its accessibility (Fig. 3B). This broader movement can be appreciated by plotting the distance between residue S680, placed at the tip of the loop and flanking the furin-like domain, and S940, located on the HR1 domain: this distance significantly increases in the G614 variant reaching a mean value of 2.1 ±3 nm (Fig. 3C-D).

**Figure 3.**
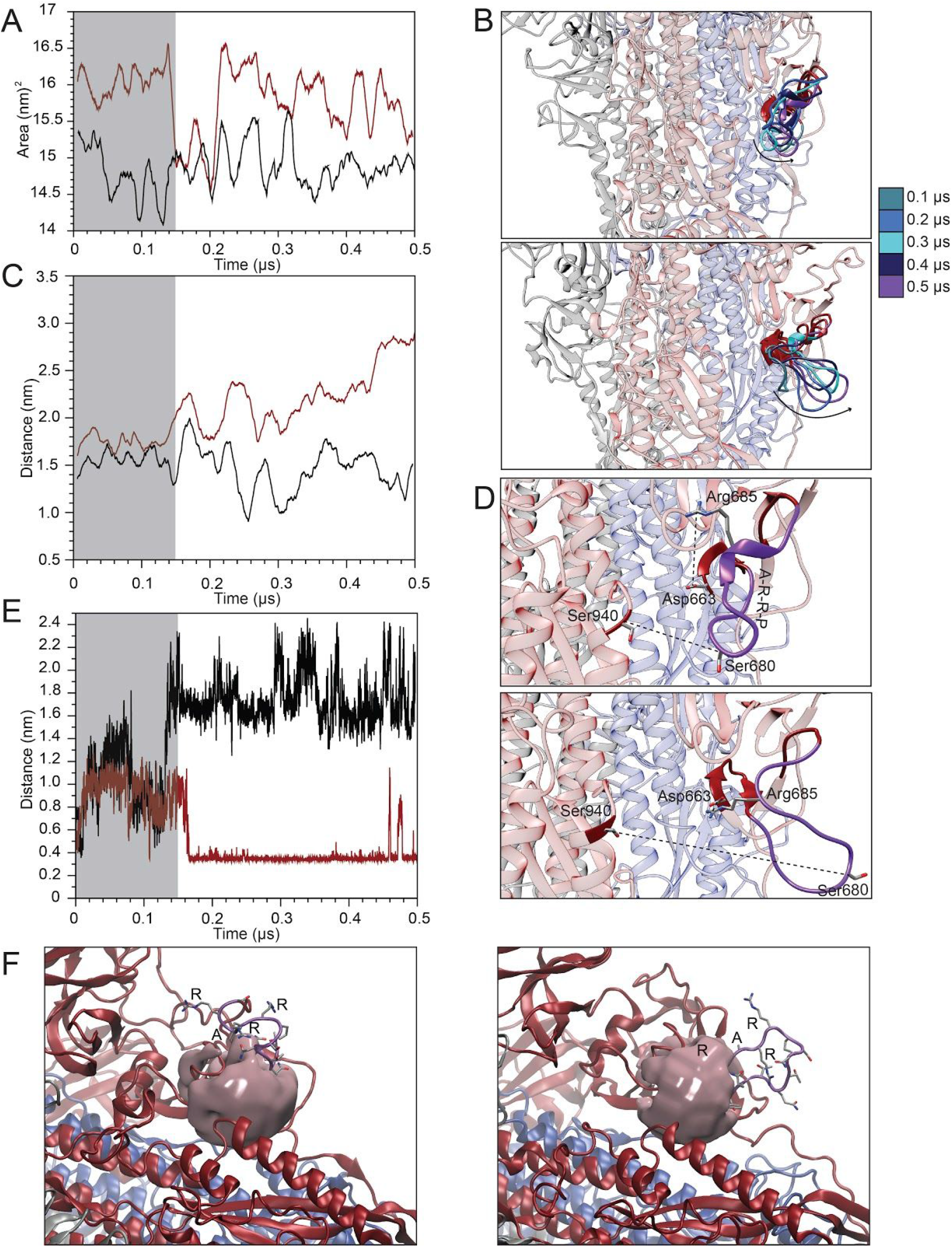
Analysis of the furin-like domain. **A.** Variation of the solvent accessible surface area of residues 675-692 in the D614 (black) and G614 (red) proteins. **B.** Superposition of the structures extracted from the D614 (upper panel) and G614 (lower panel) trajectories showing the displacement of the furin-like domain. **C.** Variation of the atomic distance calculated as a function of time between the lateral chains of S680 and S940. **D.** Representative snapshots showing the relative positions of residues S680-S940 and R685-Asp663 in structures extracted from the D614 (upper panel) and G614 (lower panel) trajectories. **E.** Variation of the atomic distance calculated as a function of time between the lateral chains of R685 and D663. **F.** Representative snapshot showing the volume of the cavity formed between the furin-like domain and the surrounding structural elements in D614 (left panel) and G614 (right panel).

In addition, as a consequence of this displacement of the loop, R685, the residue of the furin-like domain directly targeted by the protease is fixed by an electrostatic interaction formed with D663 (Fig. 3D). This salt bridge is present in the G614 variant and detected for 70% of the time, while it is never observed in the D614 variant (Fig. 3E). Finally, we detect an increase in the volume of the cavity formed by the multibasic loop and the surrounding structural elements. At 0.5 µs of the simulation (i.e. at the end of the simulation) we measured a volume of 1.46 nm^3^ and 1.7 nm^3^ for the D614 and G614 variants respectively (Fig. 3F). The opening of the loop in the furin-like domain, indeed, increases the size of the channel at the interface with the HR1 domain where the protease can be accommodated.

Taken together, these data indicate that the presence of a glycine in position 614 has a direct effect on the dynamics of the furin-like domain when the RBD is in the active up conformation (*i.e.*, in the active state). The increased mobility of the loop harboring the multibasic proteolytic site increases the accessibility of the furin-like domain to the solvent and fixes the position of R686. This likely facilitates the site recognition by the protease and promotes the subsequent cleavage, leading to an increased rate of spike protein priming. As the furin-like cleavage site an evolutionary novelty in SARS-CoV-2 as compared to its close CoVs relative (Andersen et al 2020, Xiao et al 2020), and given its crucial role for efficient cell-cell transmission (Hoffman et al 2020a and 2020b), our results argue for a refined efficiency of this infection mechanism in the G614 variant.

### The D614G substitution has a long-range effect on the RBD dynamics

To filter out the noise given by minor movements that could hamper the analyses of the main motions that dictate the large conformational movement of a protein/domain (D’Annessa et al 2018, 2019), we decomposed the whole protein motion in its principal components (PCA) and discuss the most relevant motions described by the first three eigenvectors. The two variants of the spike display clear differences in the amplitude of the conformational space sampled by the RBD in the up conformation along the first three eigenvectors (Fig. 4A and Fig. S3), together describing more than 50% of the total motion, whereas the essential dynamics is largely similar between the two variants for the chains with the RBD in down conformation (data not shown). In general, the motion of the whole S1 subdomain, including the RBD, is more spread in D614, while the same region in G614 has a more confined motion. However, the receptor-binding motif (RBM residues 435-506), the apical region of the RBD directly deputed to bind the ACE2 receptor, shows a larger movement, also allowing it to move far apart from the NTD, as evidenced by the increase in the distance between these two domains in G614 (Fig. 4B). Conversely, even if the NTD and RBD in D614 appear to sample a larger space, their relative position is fixed along the simulation (Fig. 4B).

**Figure 4.**
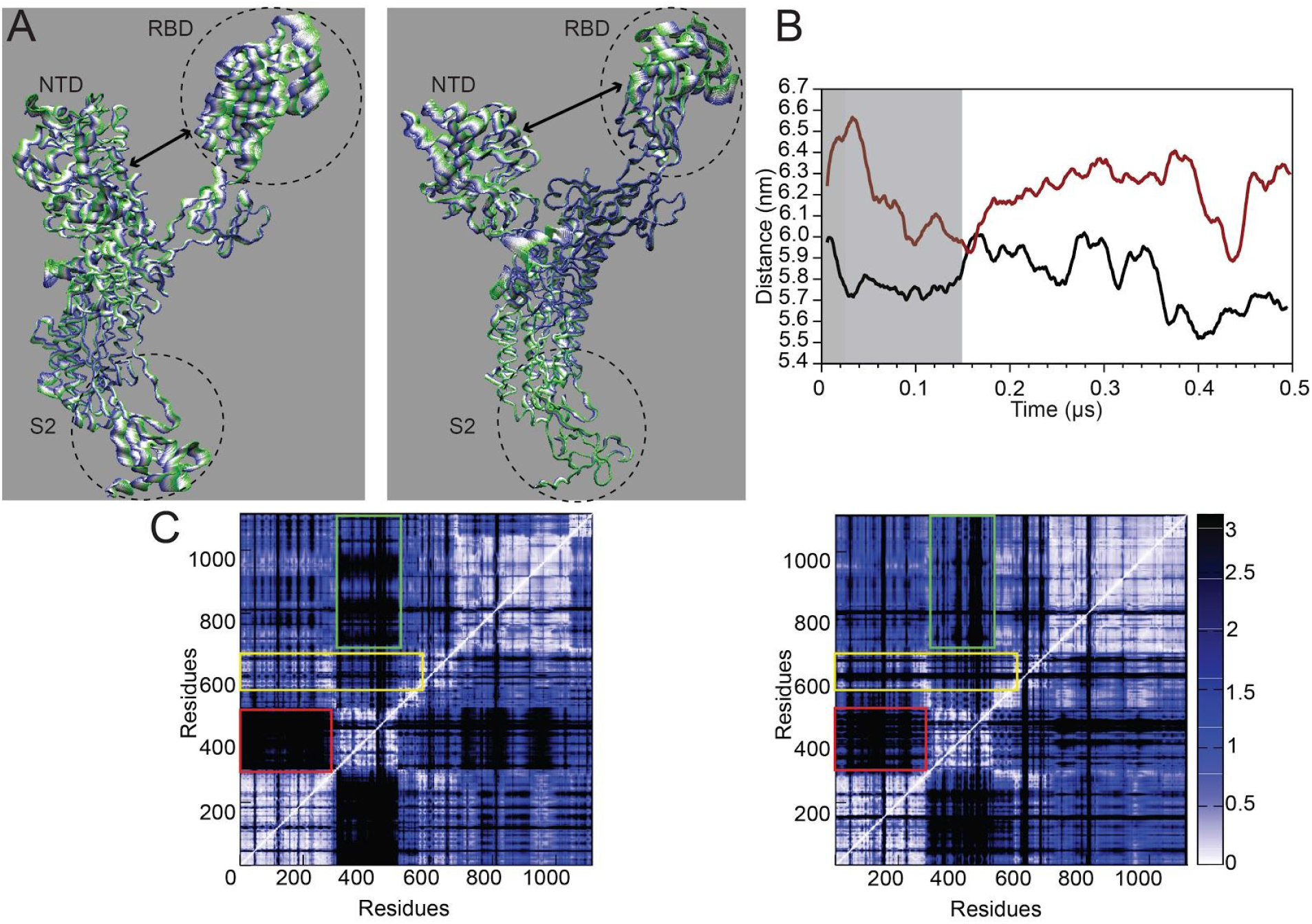
Principal component analysis of chain A. A) Projection of the motion along the first eigenvector for the chain A of D614 (left) and G614 (right). The color code from green to blues and the thickness of the tube identifies the amplitude of the motion. B) Evolution of the distance calculated between the center of mass of the NTD and the RBD in D614 (black) and G614 (red) as a function of time. C) Distance fluctuation matrices calculated for chain A in D614 (left) and G614 (right). The rectangles underline the regions with differences between the two proteins.

The differences in the inter-domains coordination can be better highlighted by plotting the matrix reporting the distance fluctuation (DF) between pairs or residues (Fig. 4C). In this way we can filter out the regions that are connected dynamically and functionally by following the variation in the relative distance among selected residues (Morra et al 2012, D’Annessa et al 2019). Basically, a variation in the pairs distance is an index of low dynamical connection between the two residues in the pair. In D614 the dynamical connection between the NTD and the RBD is low (Fig. 4C, left panel, red rectangle), as well as that between the RBD and the S2 domain (Fig. 4C, left panel, green rectangle). On the contrary, the NTD-RBD connection becomes stronger in the G614 variant, once again evidencing that the two domains are dynamically and functionally more connected (Fig. 4C, right panel, red rectangle). In this case, a very strong connection also appears between the RBD and the S2 domain (Fig. 4C, right panel, green rectangle), highlighting a stronger functional inter-domains connection in the G614 variant. Interestingly, in G614 residues 675-692 are poorly connected with the rest of the protein, when compared to D614 (Fig. 4C, yellow rectangles), once again confirming the peculiar motion of the furin-like domain in the G614 variant.

The dynamics of the RBD domain is therefore different in the two 614 variants, with the domain in G614 exploring a more open conformation, suggesting a long range effect of the D to G substitution that likely increases the efficient interaction of RBD with the ACE2 receptor. This is in line with the recent cryo-EM structure of the cognate SARS-Cov spike protein bound to ACE2 provided by Song et al. (Song et al. 2018) where the ACE2-bound RBD is shown as more open when compared to the ACE2-free RBD in the up conformation. This increased opening of the RBD domain with respect to the vertical axis of the trimer is a prerequisite for allowing the proper interaction with the receptor and for further transition from pre- to post-fusion conformation.

## CONCLUSIONS

The G614 spike variant rose to high frequency in the genomic samples from the vast majority of countries in the world (Korber et al 2020). By statistically controlling for random genetic drift within geographical areas and for migration among different areas, we confirmed that the available data are compatible with a selective advantage of G614. Moreover, the increasing frequencies in the genomic samples are mirrored by similar trends in the actual virus populations reconstructed by a coalescent-based approach, which argues against a sampling bias in favor of G614 due to enhanced severity of the COVID-19 symptoms.

The structural analysis of the spike protein revealed the crucial role of position 614 in interacting with both the S1/S2 furin-like and the receptor-binding domain. The substitution in this position of an aspartic acid (D), carrying a negatively charged side chain, with a glycine (G) strongly affects the dynamics of the spike functional domains at a short- and long-range, making it sample novel structural conformations, responsible for the higher fitness of this SARS-Cov-2 variant. In particular, the long-range effect of the D614G substitution influences the internal dynamics of the RBD and its relative orientation, allowing it to acquire a more open conformation which is particularly suitable for the interaction with the host ACE2 receptor. Nevertheless, the stronger dynamical connection of the RBD with the S2 domain could allow to remove the steric restraints on helix linker 2, which would better trigger the release of the S1 subunits, allowing the extension of pre-fusion S2 helixes to form the postfusion S2 long helix bundle (Song et al 2018).

Finally, compared with the ancestral form D614, the G614 variant shows a marked conformational plasticity of the furin-like domain, also increasing the volume of the cavity surrounding the cleavage site. These two structural and dynamical features possibly favor the furin-like domain recognition by the protease and improve the efficiency of the proteolytic cleavage, a crucial step for host cell infection. However, what seems to be the evolutionary strength of the virus in terms of invasiveness can be transformed in its weakness, in the fight against COVID-19. Indeed, the peculiar properties of the furin-like domain, as the widening of the cavity, can be exploited to trigger the rational design of drug molecules able to bind it and compete for the interaction of the spike with the protease, thus specifically blocking the invasiveness of this highly contagious SARS-CoV-2 variant.

## Methods

### Models of temporal variation of G614 in the genomic samples

We tested whether the frequency increase of the G614 spike protein variant can be attributed to random genetic drift by fitting a generalized mixed model (GLMM) with binomial error structure. The allele at genomic position 23,403 (G/A), corresponding to amino acid substitution D → G in residue 614 of the SARS-CoV-2 spike protein, was the binomial response. The fixed effect was the sampling day. We included random intercepts and slopes for “territories” (countries, except for USA states and Chinese provinces) and a covariance structure represented by *F*_ST_ values between viral samples from each pair of territories. Our model structure assumes that random slopes of sampling day within “territories” captures the random component of allele frequency change (drift) within each territory and that *F*_ST_ values capture the relative amount of drift that is shared between pairs of territories (due to migration, *i.e.* exchange of viral lineages). We fitted the GLMM using the function fitme from *R* package “spaMM” v3.2.0 (Rousset & Ferdy, 2014) on data from territories with at least 30 sequences (*N* territories = 24, *N* sequences = 5,773). *F*_ST_ values were calculated using the function pairwise.fst (*R* package “hierfstat” v0.04-22, Goudet, 2005) and including all sequences available for each territory. Note that the latter choice is conservative, as a common increase in G614 or D614 variants due to selection in two populations would also result in a lower distance between them compared to others.

### Coalescent-based inference of A/G614 variants diffusion in the population

Complete high-coverage whole genome sequences of SARS-CoV-2 from European, US, Australian and Chinese isolates were downloaded from the GISAID EpiFlu™ Database on 24 Apr and 6 May 2020. A total of 6,655 sequences were aligned to the Wuhan-Hu-1 SARS-CoV-2 reference sequence (MN908947.3; Wu et al 2020) using mafft (Katoh and Toh 2008). Alignment was checked by eye in Aliview (Larsson 2014) to remove indels, and then parsed using a custom *python* script to trim sequences before position 150 and after position 29150, and discard sequences with more than 500 missing bases.

Sequences were grouped according to the nucleotidic allele at position 23,403 (whether A or G, corresponding to the D614 and G614 variants, respectively) and then by territory, resulting in two alignments per territory. Only pairs of alignments with at least 30 sequences for each of the two alleles were retained for downstream analyses (Australia, Belgium, England, Iceland, Netherlands, Spain, USA-NY, USA-WA, USA-WI, and Wales). The maximum number of sequences to be used in the coalescent-based analysis was limited to 250, which were randomly selected when needed. Three independent replicates of randomly selectected sequences were run and checked for consistency of the results.

The demographic history of the D/G614 SARS-CoV-2 variants in each territory was reconstructed using the Bayesian Skyline plot analyses as implemented in Beast v2.6 (Bouckaert et al 2019). For each alignment we prepared the input file by setting the tips dates as days before the most recent sequence (available as “Collection date” in the GISAID metadata), a HKY substitution model, a strict clock model, a coalescent Bayesian Skyline as tree model and 100,000,000 iterations for the MCMC chain. We ran three replicates for each alignment and checked their convergence in Tracer 1.8 (Rambaut et al 2018). Bayesian Skyline plot analyses (Drummond et al 2005) were run in Tracer and results exported as tabular values. We then calculated the relative frequency through time of the G614 variant in the population by dividing the median values of the estimated effective population size of the G614 by the sum of the median values of the estimated effective population sizes of A and G614 (*N*_e_(A) and *N*_e_(G), respectively). The confidence intervals of the estimated relative frequency of the G614 were calculated as follows: for the upper boundary = 97.5% *N*_e_(G) / (97.5% *N*_e_(G) + 2.5% *N*_e_(A)); lower boundary = 2.5% *N*_e_(G) / (2.5% *N*_e_(G) + 97.5% *N*_e_(A)). Plots were drawn in *R* and *python* using standard plotting functions and libraries.

### Molecular Dynamics Simulations of D614 and G614 variants

The starting structure of the Spike trimeric complex with one RBD in up and two in down conformation was taken from the structure deposited in the protein data bank with code 6VSB (Wrapp et al 2020). Missing residues, mainly belonging to loops regions, were reconstructed using the SwissModel web server (https://swissmodel.expasy.org/). Because of the lack of a reliable template to model the 3D arrangement of the HR2 segment with respect to the rest of the protein, and thus to model the transmembrane region, the model covers residue 27-1146. All the N-acetylGlucosamine residues present in the cryo EM structure bound to asparagines residues were retained in the system. The D to G mutation in position 614 was introduced with the Chimera program (Pettersen et al 2004) and the structure obtained was further minimized. Topologies of the two systems were built using tleap with the AMBER14 force field (Case at al 2014). Each protein was then placed in a triclinic simulative box filled with TIP3P water molecules (Jorgensen et al 1983). Addition of sodium counterions rendered the systems electroneutral; each system consisted of ∼555.000 atoms.

The simulations were carried out with amber14 using pmemd.CUDA (Case et al 2014). The systems were first minimized with 10000 steps of steepest descent followed by 10000 steps of conjugate gradient. Relaxation of water molecules and thermalization of the system in NPT environment were run for 1.2 ns at 1 fs time-step. In detail, 6 runs of 200 ps each were carried out by increasing the temperature of 50 K at each step, starting from 50 K to 300 K. The systems were then simulated with a 2 fs time-step for 500ns each in periodic boundary conditions, using a cut-off of 8 Å for the evaluation of short-range non-bonded interactions and the Particle Mesh Ewald method (Cheatham et al 1995) for the long-range electrostatic interactions. The temperature was kept constant at 300 K with Langevin dynamics (Ceriotti et al 2009) and pressure fixed at 1 Atmosphere through the Langevin piston method (Feler et al 1995). The bond lengths of solute and water molecules were restrained with the SHAKE Ryckaert et al 1977) and SETTLE algorithms (Miyamoto and Kollman 1992), respectively. As stated before, the transmembrane region is lacking. In order to mimic the binding of the spike trimer on the viral membrane, we applied a force of 1000 kj on the last four residues (1143-1146) to anchor the protein. Analyses were carried out using Gromacs 5 package (Hess et al 2008) or with VMD (Humphrey et al 1996) and custom code.

As reported in supplementary figure S2 (panels C and D), both proteins largely deviate from their starting conformation, reaching a Root Mean Square Deviation (RMSD) value of ∼0.4 nm. This was however expected and due to the fact that the starting configuration used to carry out the simulations comes from cryoEM and was solved at a resolution of 3.46 Å. It is then reasonable that once hydrated, the proteins undergo sudden conformational changes to relax the structure. However, the RMSD in both cases reaches a plateau at around 0.15 µs, meaning that the two proteins find the stability, and for this reason all analyses reported here have been performed on the last 0.35 µs of simulative time.

Volumes were computed using POVME (Wagner et al 2017). An inclusion sphere with a radius of 11 Å was manually placed between the furinic loop and the near S2 domain after careful optimization in VMD (Humphrey et al 1996).

Principal component analysis (PCA) for the bound and unbound trajectories was carried out on the 3N×3N Cartesian displacement matrix whose elements are calculated as:

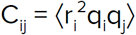

N being the number of Cα atoms and q_i_ the (mass-weighted) displacement of the i-th Cα atoms from the reference value (after removal of rotational and translational degrees of freedom).

The matrix of the distance fluctuations DF was computed as:

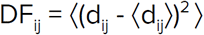

being *d*_*ij*_ the (time-dependent) distance of the *Cα* atoms between amino acids *i* and *j* and ⟨ ⟩ the time-average over the trajectory. DF is independent upon translations and rotations of the molecules and thus on the choice of a protein reference structure.

## Acknowledgements

All SARS-CoV-2 genome sequences used in this study were downloaded from the GISAID EpiFlu™ Database, www.gisaid.org (accessed on 24/04/2020 and 06/05/2020, selected locations: Europe, US, Australia, China; selected sampling date: before 13/04/20). We thank Prof. Giorgio Colombo for providing the code for the calculation of the distance fluctuation matrices.

## Supplementary Figures

**Figure S1.**
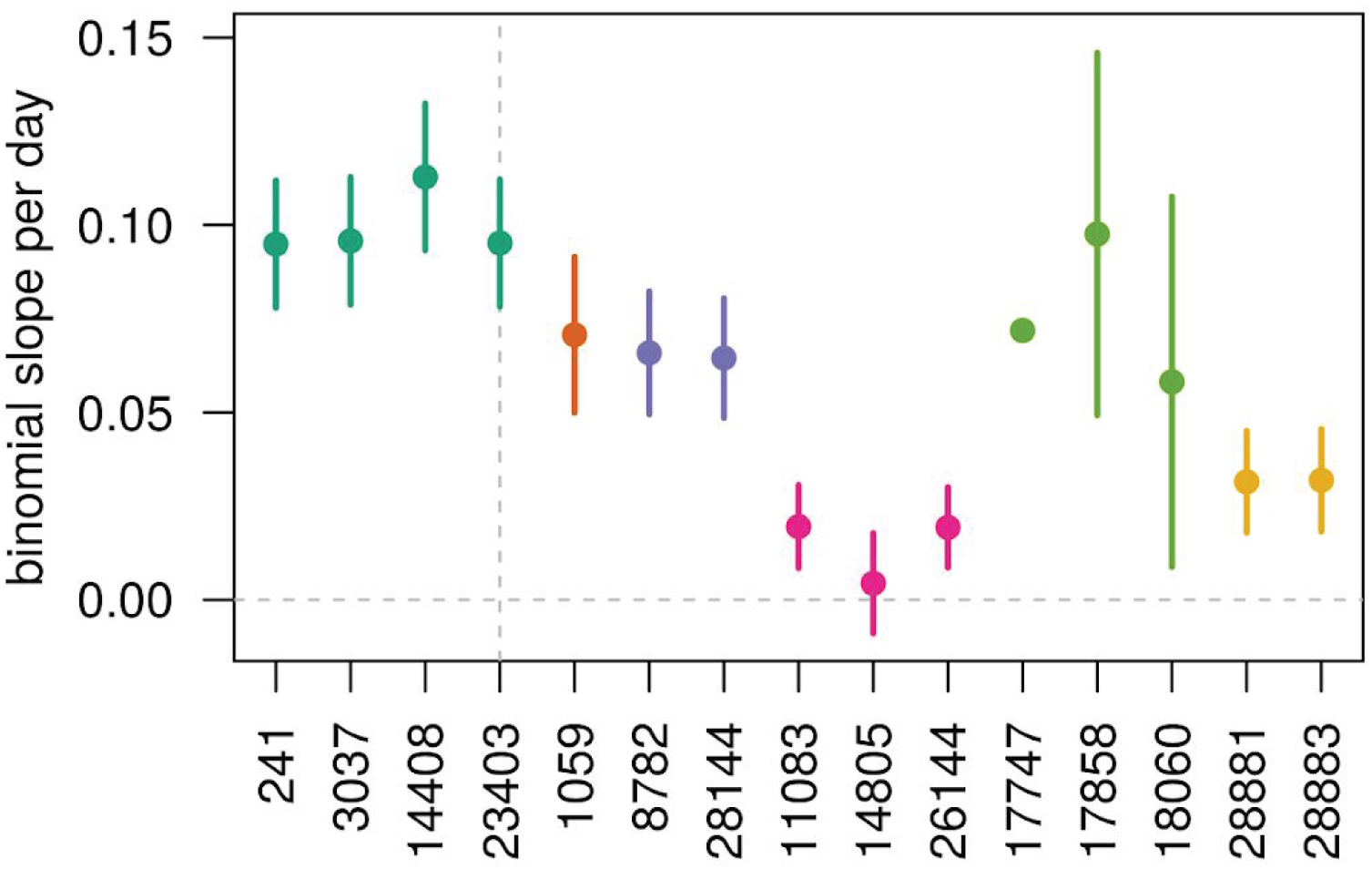
GLMM slopes (±SE) for 14 variable positions in the SARS-Cov2 genome. D/G614 is pos. 23,403 (vertical dashed line). Color by linkage group (threshold *rho* = 0.8).

**Figure S2.**
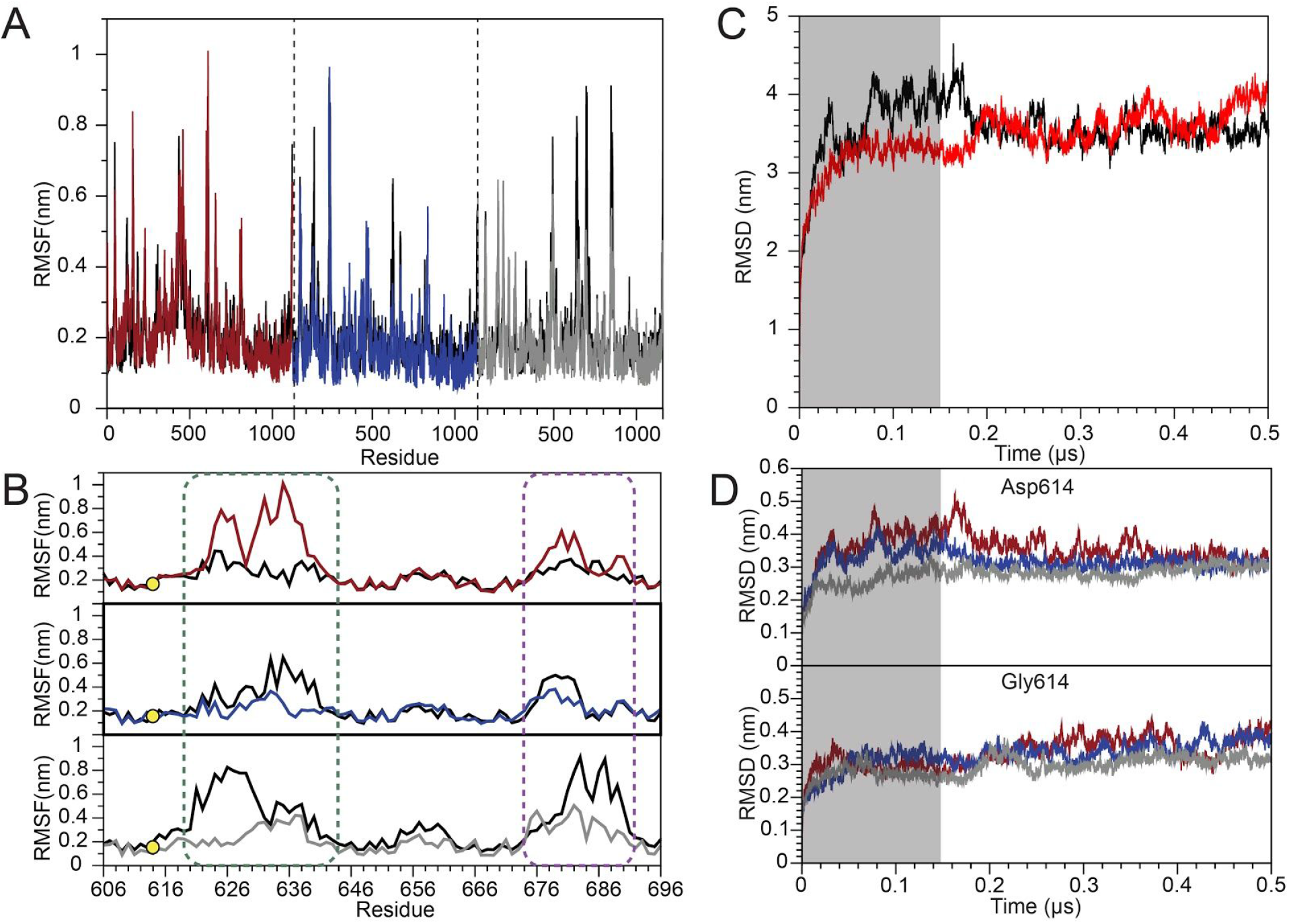
Time evolution of the systems and main fluctuations. **A.** per-residue Root Mean Square Fluctuation (RMSF) of D614 (black line) and G614 (chain A 4 red; chain B blue; chain C grey.) **B.** per-residue RMSF of residues 606-696 chain A, D614 black and G614 red. Chain B, D614 black and G614 blue. Chain C, D614 black and G614 grey. **C.** Root Mean Square Deviation (RMSD) of D614 (black line) and G614 (red line) whole trimers. The grey box highlights the part of the trajectories avoided from the analysis. **D.** RMSD of the each chain in the two proteins. Color code: chain A, red; chains B, blue; C, grey.

**Figure S3.**
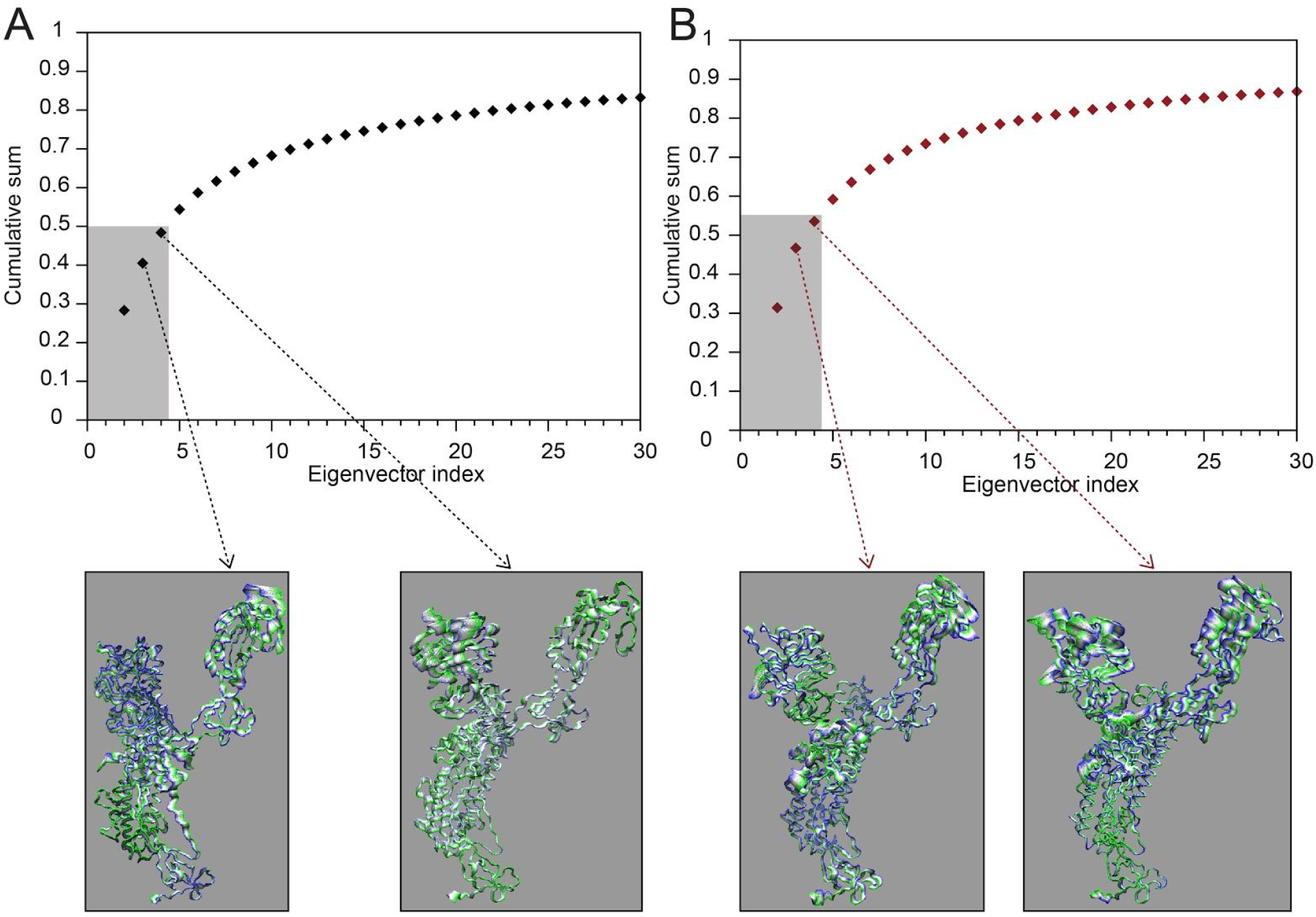
Essential Dynamics. Cumulative percentage of the weight of the single eigenvectors on the total motion of chain A in D614 **A** and G614 **B**. For the sake of clarity, both plots report only the cumulative sum of the first 30 eigenvectors. The lower panels show the projection of the motion along the second and third eigenvector for D614 (A) and G614 (B). The grey box highlights the first three eigenvectors that together describe more than 50% of total motion in both chains A and for this reason have been considered for discussion.

## References

Andersen, K.G., Rambaut, A., Lipkin, W.I., Holmes, E. C., Garry, R.F. (2020). The proximal origin of SARS-CoV-2. Nature medicine, 26(4), 450–452.

Becerra-Flores, M., Cardozo, T. (2020). SARS-CoV-2 viral spike G614 mutation exhibits higher case fatality rate. The Journal of clinical Practice, 0–2.

Berry, J.D., Jones, S., Drebot, M.A., Andonov, A., Sabara, M., Yuan, X.Y., et al. (2004). Development and characterisation of neutralising monoclonal antibody to the SARS-coronavirus. Journal of virological methods, 120(1), 87–96.

Belouzard S., Millet J.K., Licitra B.N., Whittaker G.R. (2012). Mechanisms of coronavirus cell entry mediated by the viral spike protein. Viruses.

Bouckaert R., Vaughan T.G., Barido-Sottani J., Duchêne S., Fourment M., Gavryushkina A., et al. (2019). BEAST 2.5: An advanced software platform for Bayesian evolutionary analysis. PLoS Comput Biol 15(4): e1006650.

Brufsky, A. (2020). Distinct Viral Clades of SARS-CoV-2: Implications for Modeling of Viral Spread. Journal of Medical Virology, 0-1.

Case, D.A., Babin, V., Berryman, J.T., Betz, R.M., Cai, Q., Cerutti, D.S., Cheatham, T.E.III, Darden, T.A., Duke, R.E., Gohlke, H., Goetz, A.W., Gusarov, S., Homeyer, N., Janowski, P., Kaus, J., Kolossváry, I., Kovalenko, A., Lee, T.S., LeGrand, S., Kollman P.A., et al. (2014). AMBER 14, University of California, San Francisco.

Ceriotti, M., Bussi, G., & Parrinello, M. (2009). Langevin equation with colored noise for constant-temperature molecular dynamics simulations. Physical review letters, 102(2), 020601.

Cheatham, T.I., Miller, J.L., Fox, T., Darden, T.A., Kollman, P.A. (1995). Molecular dynamics simulations on solvated biomolecular systems: the particle mesh Ewald method leads to stable trajectories of DNA, RNA, and proteins. Journal of the American Chemical Society, 117(14), 4193–4194.

Chiara, M., Horner, D.S., Gissi, C., Pesole, G. (2020). Comparative genomics suggests limited variability and similar evolutionary patterns between major clades of SARS-Cov-2. bioRxiv.

Coutard, B., Valle, C., de Lamballerie, X., et al. (2020). The spike glycoprotein of the new coronavirus 2019-nCoV contains a furin-like cleavage site absent in CoV of the same clade. Antiviral Research, 176, 104742.

D’Annessa, I., Raniolo, S., Limongelli, V., Di Marino, D., Colombo, G. (2019). Ligand Binding, Unbinding, and Allosteric Effects: Deciphering Small-Molecule Modulation of HSP90. J Chem Theory Comput, 15(11), 6368–6381.

D’Annessa, I., Gandaglia, A., Brivio, E., Stefanelli, G., Frasca, A., Landsberger, N., Di Marino, D. (2018). Tyr120Asp mutation alters domain flexibility and dynamics of MeCP2 DNA binding domain leading to impaired DNA interaction: Atomistic characterization of a Rett syndrome causing mutation. Biochim Biophys Acta Gen Subj, 1862(5), 1180–1189.

D’Arienzo, M., Coniglio, A. (2020). Assessment of the SARS-CoV-2 basic reproduction number, R0, based on the early phase of COVID-19 outbreak in Italy. Biosafety and Health.

Domingo, E., Sheldon, J., Perales, C. (2012). Viral quasispecies evolution. Microbiol. Mol. Biol. Rev., 76(2), 159–216.

Drummond, A.J., Rambaut, A., Shapiro, B.E.T.H., Pybus, O.G. (2005). Bayesian coalescent inference of past population dynamics from molecular sequences. Molecular biology and evolution, 22(5), 1185–1192.

Feller, S.E., Zhang, Y., Pastor, R.W., Brooks, B.R. (1995). Constant pressure molecular dynamics simulation: the Langevin piston method. The Journal of chemical physics, 103(11), 4613–4621.

Goudet, J. (2005). Hierfstat, a package for R to compute and test hierarchical F-statistics. Molecular Ecology Notes, 5(1), 184–186.

Hess, B., Kutzner, C., Van Der Spoel, D., Lindahl, E. (2008). GROMACS 4: algorithms for highly efficient, load-balanced, and scalable molecular simulation. Journal of chemical theory and computation, 4(3), 435–447.

Hoffmann M., Kleine-Weber H., Schroeder S., et al. (2020a). SARS-CoV-2 Cell Entry Depends on ACE2 and TMPRSS2 and Is Blocked by a Clinically Proven Protease Inhibitor. Cell, 181, 271-280.e8.

Hoffmann M., Kleine-Weber H., Pöhlmann S. (2020b). A multibasic cleavage site in the spike protein of SARS-CoV-2 is essential for infection of human lung cells. Cell Press, 78, 1–6.

Humphrey, W., Dalke, A., Schulten, K. (1996). VMD: visual molecular dynamics. Journal of molecular graphics, 14(1), 33–38.

Jorgensen, W.L., Chandrasekhar, J., Madura, J.D., Impey, R.W., Klein, M.L. (1983). Comparison of simple potential functions for simulating liquid water. The Journal of chemical physics, 79(2), 926–935.

Katoh, K., Toh, H. (2008). Recent developments in the MAFFT multiple sequence alignment program. Briefings in bioinformatics, 9(4), 286–298.

Korber, B., Fischer, W., Gnanakaran, S.G., Yoon, H., Theiler, J., Abfalterer, W., Partridge, D.G., et al. (2020). Spike mutation pipeline reveals the emergence of a more transmissible form of SARS-CoV-2. bioRxiv.

Laha, S., Chakraborty, J., Das, S. et al. (2020). Characterizations of SARS-CoV-2 mutational profile. bioRxiv, 9860, 102–109.

Larsson, A. (2014). AliView: a fast and lightweight alignment viewer and editor for large datasets. Bioinformatics, 30(22), 3276–3278.

Li, F. (2016). Structure, Function, and Evolution of Coronavirus Spike Proteins. Annual Review of Virology.

Liu, S., Xiao, G., Chen, Y., He, Y., Niu, J., Escalante, C. R., Jiang, S., et al. (2004). Interaction between heptad repeat 1 and 2 regions in spike protein of SARS-associated coronavirus: implications for virus fusogenic mechanism and identification of fusion inhibitors. The Lancet, 363(9413), 938–947.

Liu, Y., Gayle, A.A., Wilder-Smith, A., & Rocklöv, J. (2020). The reproductive number of COVID-19 is higher compared to SARS coronavirus. Journal of travel medicine.

Miyamoto, S., Kollman, P.A. (1992). Settle: An analytical version of the SHAKE and RATTLE algorithm for rigid water models. Journal of computational chemistry, 13(8), 952–962.

Morra, G., Potestio, R., Micheletti, C., Colombo, G. (2012). Corresponding Functional Dynamics across the Hsp90 Chaperone Family: Insights from a Multiscale Analysis of MD Simulations. Plos Comput Biol, 8 (3), e1002433

Mothes, W., Sherer, N.M., Jin, J., Zhong, P. (2010). Virus Cell-to-Cell Transmission. Journal of Virology, 84, 8360–8368.

Ou, X., Liu, Y., Lei, X., Li, P., Mi, D., Ren, L., Xiang, Z., et al. (2020). Characterization of spike glycoprotein of SARS-CoV-2 on virus entry and its immune cross-reactivity with SARS-CoV. Nature communications, 11(1), 1–12.

Pachetti, M., Marini, B., Benedetti, F., Giudici, F., Mauro, E., Storici, P., Zella, D., et al. (2020). Emerging SARS-CoV-2 mutation hot spots include a novel RNA-dependent-RNA polymerase variant. Journal of Translational Medicine, 18(1), 1–9.

Pak, J.E., Sharon, C., Satkunarajah, M., Auperin, T.C., Cameron, C.M., Kelvin, D.J., Rini, J.M., et al. (2009). Structural insights into immune recognition of the severe acute respiratory syndrome coronavirus S protein receptor binding domain. Journal of molecular biology, 388(4), 815–823.

Pettersen, E.F., Goddard, T.D., Huang, C.C., Couch, G.S., Greenblatt, D.M., Meng, E.C., Ferrin, T.E. (2004). UCSF Chimera—a visualization system for exploratory research and analysis. Journal of computational chemistry, 25(13), 1605–1612.

Rambaut, A., Drummond, A.J., Xie, D., Baele, G. and Suchard, M.A. (2018). Posterior summarisation in Bayesian phylogenetics using Tracer 1.7. Systematic Biology. Syy032.

Rousset, F., & Ferdy, J.B. (2014). Testing environmental and genetic effects in the presence of spatial autocorrelation. Ecography, 37(8), 781–790.

Ryckaert, J.P., Ciccotti, G., Berendsen, H.J. (1977). Numerical integration of the cartesian equations of motion of a system with constraints: molecular dynamics of n-alkanes. Journal of computational physics, 23(3), 327–341.

Qian, Z., Dominguez, S.R., Holmes, K.V. (2013). Role of the spike glycoprotein of human Middle East respiratory syndrome coronavirus (MERS-CoV) in virus entry and syncytia formation. PloS one, 8(10).

Simmons, G., Zmora, P., Gierer, S., Heurich, A., Pöhlmann, S. (2013). Proteolytic activation of the SARS-coronavirus spike protein: Cutting enzymes at the cutting edge of antiviral research. Antiviral Research, 100, 605–614.

Song, W., Gui, M., Wang, X., Xiang, Y. (2018). Cryo-EM structure of the SARS coronavirus spike glycoprotein in complex with its host cell receptor ACE2. PLoS Pathogens, 14, 1–19.

Vasilarou, M., Alachiotis, N., Garefalaki, J., Beloukas, A., Pavlidis, P. (2020) Population genomics insights into the recent evolution of SARS-CoV-2. bioRxiv

Walls, A.C., Tortorici, M.A., Snijder, J., Xiong, X., Bosch, B. J., Rey, F. A., Veesler, D. (2017). Tectonic conformational changes of a coronavirus spike glycoprotein promote membrane fusion. Proceedings of the National Academy of Sciences, 114(42), 11157–11162.

Walls, A.C., Park, Y.J., Tortorici, M.A. et al. (2020). Structure, Function, and Antigenicity of the SARS-CoV-2 Spike Glycoprotein. Cell, 181, 281-292.e6.

Wan, Y., Shang, J., Sun, S., Tai, W., Chen, J., Geng, Q., Zhou, Y., et al. (2020). Molecular mechanism for antibody-dependent enhancement of coronavirus entry. Journal of virology, 94(5).

Wang, S.F., Tseng, S.P., Yen, C.H., Yang, J.Y., Tsao, C.H., Shen, C.W., Huang, J.C. (2014). Antibody-dependent SARS coronavirus infection is mediated by antibodies against spike proteins. Biochemical and biophysical research communications, 451(2), 208–214.

Wagner, J.R., Sørensen, J., Hensley, N., Wong, C., Zhu, C., Perison, T., Amaro, R.E. (2017) POVME 3.0: Software for Mapping Binding Pocket Flexibility. J. Chem. Theory Comput, 13(9), 4584–4592

White, J.M., Whittaker, G.R. (2016). Fusion of Enveloped Viruses in Endosomes. Traffic, 17, 593–614.

WHO (2020a) Coronavirus disease 2019 (COVID-19) Situation Report 51. https://www.who.int/docs/default-source/coronaviruse/situation-reports/20200311-sitrep-51-covid-19.pdf?sfvrsn=1ba62e57_10

WHO (2020b) Coronavirus disease 2019 (COVID-19) Situation Report 112. chrome-extension://oemmndcbldboiebfnladdacbdfmadadm/ https://www.who.int/docs/default-source/coronaviruse/situation-reports/20200511-covid-19-sitrep-112.pdf?sfvrsn=813f2669_2.

Wrapp, D., Wang, N., Corbett, K.S., et al. (2020). Cryo-EM structure of the 2019-nCoV spike in the prefusion conformation. Science, 367, 1260–1263.

Wu, F., Zhao, S., Yu, B., Chen, Y.M., Wang, W., Song, Z.G., Yuan, M.L., et al. (2020). A new coronavirus associated with human respiratory disease in China. Nature, 579(7798), 265–269.

Xiao, K., Zhai, J., Feng, Y., et al. (2020). Isolation of SARS-CoV-2-related coronavirus from Malayan pangolins. Nature https://doi.org/10.1038/s41586-020-2313-x

Zhang, T., Wu, Q., Zhang, Z. (2020). Probable Pangolin Origin of SARSCoV-2 Associated with the COVID-19 Outbreak. Curr. Biol. 30, 1346–1351.e2.

Zhou, P., Yang, X.L., Wang, X.G., Hu, B., Zhang, L., Zhang, W., Chen, H. D., et al. (2020). A pneumonia outbreak associated with a new coronavirus of probable bat origin. Nature, 579(7798), 270–273.

